# Metabolite Relaxation Times at 3T From Birth to 7 Years

**DOI:** 10.64898/2026.10.02.754450

**Authors:** Saipavitra Murali-Manohar, Abdelrahman Gad, Zahra Shams, Gizeaddis L. Simegn, Yulu Song, Aaron T. Gudmundson, Helge J. Zo□llner, Christopher W. Davies-Jenkins, Dunja Simicic, Vivek Yedavalli, Borjan Gagoski, Ellen P. Grant, Douglas C. Dean, Ralph Noeske, Jessica L. Wisnowski, Mary Beth Nebel, Georg Oeltzschner, Richard A.E. Edden

## Abstract

**Purpose:** With growing interest in pediatric MRS to understand the neurochemistry of brain development, it is essential to establish pediatric-specific metabolite T_1_ and T_2_ relaxation times for reliable quantification of metabolites. Furthermore, characterization of age-related changes in relaxation times may provide insight into developmental differences in tissue microstructure and cellular composition. Therefore, the goals of this study were two-fold: (1) to estimate metabolite T_1_ and T_2_ relaxation times in a pediatric cohort; and (2) to assess linear relationships between age and relaxation times.

**Methods:** TI- and TE-series data were acquired from the bilateral thalamus in 20 and 25 participants aged between 0 and 7 years, respectively. Data were processed and modelled in Osprey 2.5.0. Metabolite T_1_ and T_2_ relaxation times of total N-acetylaspartate at 2.0 ppm (tNAA_2.0_), total creatine at 3.0 ppm (tCr_3.0_), total creatine at 3.9 ppm (tCr_3.9_), total choline (tCho), myo-inositol (mI) and the combined glutamate and glutamine (Glx) were estimated by fitting the modelled signal amplitudes to TI-series inversion recovery signal and TE-series exponential decay equations, respectively. Linear relationships with age were tested for metabolite T_1_ and T_2_ relaxation times using Spearman correlations.

**Results:** tNAA_2.0_ and Glx showed significant changes in T_1_ with age. tNAA_2.0_, tCr_3.0_, tCho and mI showed significant decrease in T_2_ with age.

**Conclusion:** Several metabolites showed significant changes in relaxation times with age in the thalamus region. Therefore, use of age-specific relaxation correction for pediatric studies using the linear equations provided is encouraged for reliable quantification of metabolites.

## Introduction

Rapid brain development in the early years of life is accompanied by dynamic and substantial changes in neurochemistry^1–3^ as well as tissue properties^4,5^. Proton magnetic resonance spectroscopy (^1^H-MRS) can measure neurometabolite levels, relying upon quantification relative to internal reference signals^6,7^. In essence, the ratio between a metabolite signal and the reference signal is proportional to the ratio between the metabolite concentration and the reference concentration. The measured signal, however, is attenuated by the acquisition-dependent relaxation weighting of the respective metabolite. Metabolite concentration levels can therefore be reliably estimated from signal ratios only after correcting for relaxation behavior. MRS studies often use the solvent water signal as an in vivo reference^6,8^, while accounting for the differing MR visibility and relaxation properties of water across tissue compartments.

MRS measurements are typically lengthy, and it is impractical to measure metabolite relaxation parameters for every subject, especially in infants and young children. Therefore, consensus practice in the area is to acquire the spectrum, model the signals therein, and then use established relaxation values from literature to estimate metabolite levels^7^. In contrast to adult MRS, where there is an extensive literature of relaxation values^9,10^, including age-dependent reference curves^11,12^, the pediatric relaxation literature is limited^13–15^. A few pediatric studies of neurometabolite trajectories have used pediatric-specific relaxation corrections^2,13,15^, although these have mostly relied upon individual measurements in the same subjects. The majority of pediatric studies have either used metabolite ratios without correcting for relaxation, the water reference signal without relaxation correction, values from a lower field strength or inappropriate adult relaxation reference values to ‘correct for’ relaxation^3,16–19^. Without a comprehensive relaxation reference literature, concentration estimates from pediatric MRS will be biased to an age-dependent degree, potentially leading to incorrect interpretation of changes in metabolite levels during development.

Measurements of metabolite T_1_ and T_2_ relaxation times themselves may act as valuable biomarkers of tissue properties. There is strong evidence that some metabolites are preferentially localized to certain cell types (e.g. NAA being more concentrated in neurons^20^), so metabolite-specific relaxation times may be sensitive to cell-type-specific changes in the cellular microenvironment. In fact, metabolite relaxation times have been shown to vary with certain pathologies. For instance, metabolite T_2_ relaxation times were observed to change in acute hypoxia-ischemia in newborn piglets, later stages of focal ischemia in rat brain, HIV patients, pediatric brain tumors^21^ and are also different in low-grade and high-grade gliomas^22–25^. Increased lactate T_2_ relaxation times along with reduced NAA levels were linked to patients with the worse outcome in perinatal cerebral hypoxic-ischemic injury^26^.

MRS is increasingly used in pediatric research, including large, multicenter NIH-funded studies like HEALthy Brain and Child Development (HBCD)^19^ and BRAINY-Type 1 Diabetes. Therefore, there is a pressing need for metabolite relaxation reference literature spanning infancy and early childhood. In this study, we report T_1_ and T_2_ relaxation times of the major metabolites in the bilateral thalamus from birth to 7 years of age at 3T. For metabolites that showed significant changes in relaxation with age, metabolite T_1_ and T_2_ relaxation models are expressed as a function of age for convenient use in other pediatric MRS studies.

## Methods

### Study Participants

25 children ages 0 to 7 years 11 months were recruited for this study at the Johns Hopkins University School of Medicine, Baltimore, MD, USA. TI- and TE-series data were collected on 20 (8 female) and 25 (12 female) participants, respectively. Written informed consent was obtained from the parent or legal guardian following site IRB guidelines. Data were acquired during natural sleep for children under the age of 2 years.

Children older than 2 years were trained to lie still during a mock scan before the MR scan and watched movies or cartoons during the scan.

### Data Acquisition

All data were acquired on a 3T Philips dStream Ingenia Elition MRI scanner using a 32-channel-receive head coil. In response to gray-matter-white-matter contrast changes early in life, voxel placement was performed on T_2_-weighted images acquired for children less than 1 year old and T_1_-weighted images for older children. A 30 (RL) × 23 × 23-mm^3^ MRS voxel was placed in the bilateral thalamus (matching the voxel of interest for the HBCD study^19^). PRESS-localized inversion recovery TI- and echo time TE-series data were acquired to model T_1_ and T_2_ relaxation times, respectively. 8 inversion times (TI: 300; 400; 511; 637; 780; 947; 1148; 1400 ms) were acquired for the TI series with TR/TE: 2000/30 ms. 10 echo times (TE: 30; 50; 74; 101; 135; 179; 241; 350; 550; 800 ms) were acquired for the TE series with TR 3s. 24 and 16 transients were acquired per TI and TE, respectively. Supplementary Table 1 provides additional acquisition details for both TI and TE series.

**Table 1.** T_1_ and T_2_ relaxation times of metabolites. Intercept values provide reference relaxation times at birth (0 months). Slope values of the linear curves from Figures 3 and 4 are provided for metabolites that show significant change in relaxation times with age. The parametrized model (Slope × age + Intercept) will provide estimated reference relaxation times for a given age in months. Mean and standard deviations (Std) of T_1_ and T_2_ relaxation times are provided for each metabolite in ms.

| Metabolites | $T_1$ | | | $T_2$ | | |
| --- | --- | --- | --- | --- | --- | --- |
| | Mean ± Std<br>ms | Intercept<br>ms | Slope<br>ms/month | Mean ± Std<br>ms | Intercept<br>ms | Slope<br>ms/month |
| tNAA<sub>2.0</sub> | 1227 ± 55 | 1179 | 0.95 | 302 ± 51 | 365 | -1.16 |
| tCr<sub>3.0</sub> | 1190 ± 68 | | | 170 ± 25 | 197 | -0.51 |
| tCr<sub>3.9</sub> | 854 ± 35 | | | 143 ± 29 | 165 | -0.39 |
| tCho | 1120 ± 51 | | | 291 ± 63 | 366 | -1.39 |
| mI | 1157 ± 107 | | | 183 ± 48 | 245 | -1.15 |
| Glx | 1328 ± 214 | 1496 | -3.27 | 257 ± 81 | | |

### Data Analysis

TE-specific metabolite basis sets were simulated in MRSCloud^27^, consisting of 19 metabolite basis functions, as well as 5 macromolecule (MM) and 3 lipid Gaussian basis functions. Supplementary Table 2 contains the full list, alongside other relevant modelling parameters for both TI- and TE-series models.

For the TI series, metabolite basis sets for TE 30 ms were tailored with TI-appropriate polarity for metabolites and macromolecules, and data acquired at TI of 637 ms were modelled with a reduced basis set and a more flexible spline baseline (knot spacing: 0.2). TI-637 and TI-780 ms were modelled without lineshape convolution. These steps, which have been applied previously^11^, accommodate the nuances of spectra at each inversion time. TI- and TE-series data were modelled in Osprey 2.5.0^28^. No datasets were excluded from T_1_ analysis. 4 datasets were excluded from T_2_ analyses due to broad linewidth and poor SNR. T_1_ and T_2_ relaxation times were then determined from secondary exponential modeling for total N-acetyl aspartate at 2.0 ppm tNAA_2.0_, total creatine at 3.0 ppm tCr_3.0_, total choline tCho, total creatine at 3.9 ppm tCr_3.9_, myo-inositol mI and the sum of glutamate and glutamine, Glx. Data for TE 800 ms were excluded from these models due to difficulties with primary linear combination modeling of the extremely low-SNR spectra. T_1_ and T_2_ exponential models with R^2^ < 0.5 were to be excluded from further analyses, as applied previously^11^. Based upon the modest sample size and visual inspection of the Q-Q plot of age values which did not support normality, Spearman rank correlation was used to assess associations between age and relaxation times. r is the Spearman correlation coefficient, when positive indicated metabolite T_1_ and T_2_ increasing with age and negative indicating those decreasing with age. Only those associations with *p* < 0.05 were considered statistically significant.

## Results

TI- and TE-series spectra from a 2-month-old infant and 7½ year-old (90-month old) child, are shown in Figures 1A and 2A respectively, along with models and fit residuals for each spectrum. These subjects were chosen to illustrate spectra from the two extremes of the cohort age range. Inversion-recovery exponential modeling to estimate T_1_ relaxation times of the 6 metabolite signals of interest for the 2-month-old infant is presented in Figure 1B. Across all subjects, T_1_ models for tNAA_2.0_, tCr_3.0_, tCho and mI had R^2^ > 0.96, indicating good fits to the datapoints. All R^2^ for tCr_3.9_ and Glx were above 0.64 and 0.75, respectively. Exponential modeling for estimating T_2_ relaxation times for the metabolite signals is presented in Figure 2B, for the same infant. T_2_ models for all metabolites in all subjects had R^2^ > 0.72, also suggesting good fits to the datapoints. There were no T_1_ or T_2_ exponential models that exceeded the R^2^ < 0.5 secondary-model criterion, and therefore none were excluded from further analysis.

**Figure 1:**
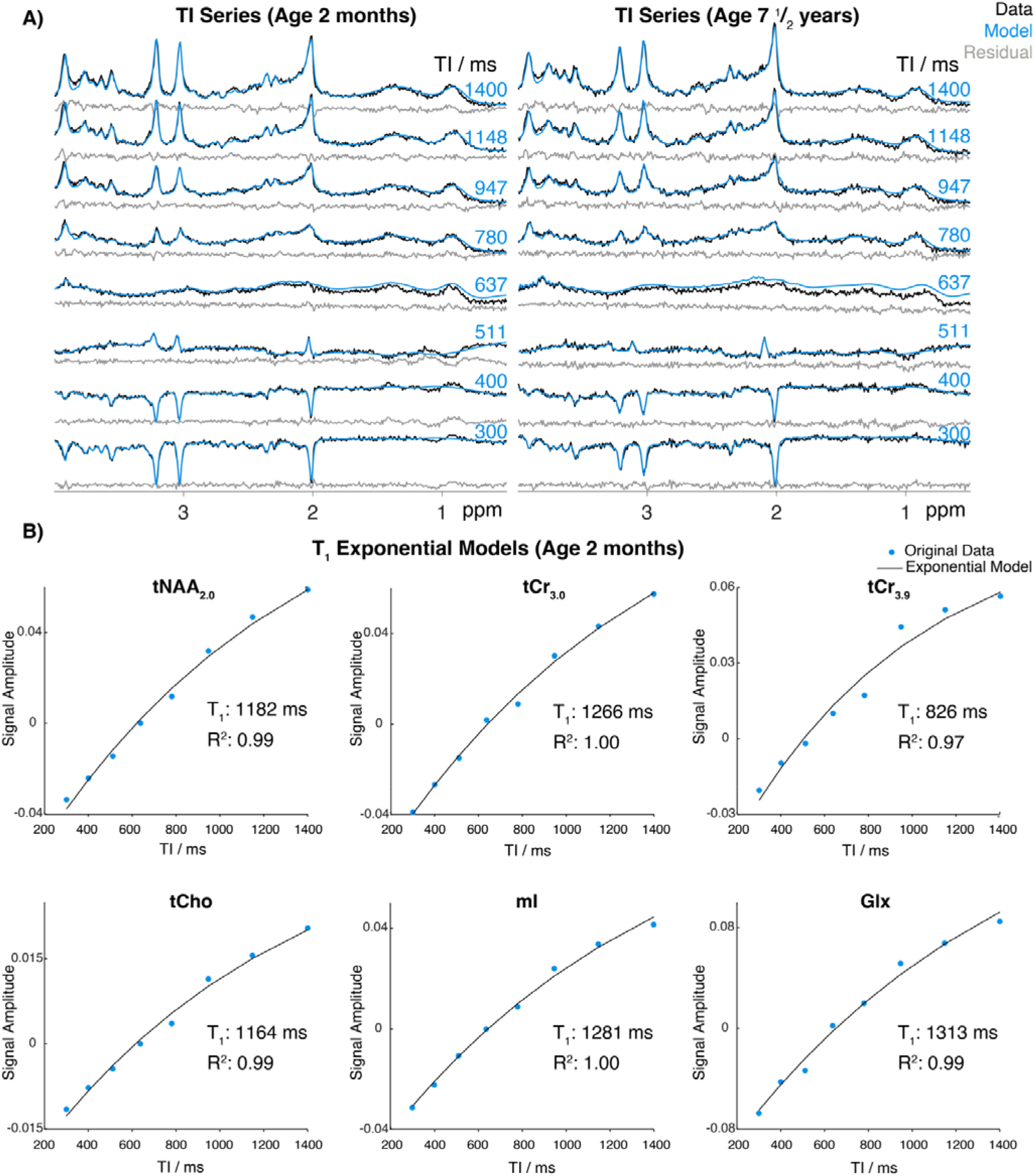
**A)** Example TI-series spectra, models and fit residuals from a 2-month-old infant and 90-month-old child. B) Inversion-recovery modeling to determine metabolite T_1_ relaxation times, using modelled metabolite coefficients from the 2-month-old spectra in A.

**Figure 2:**
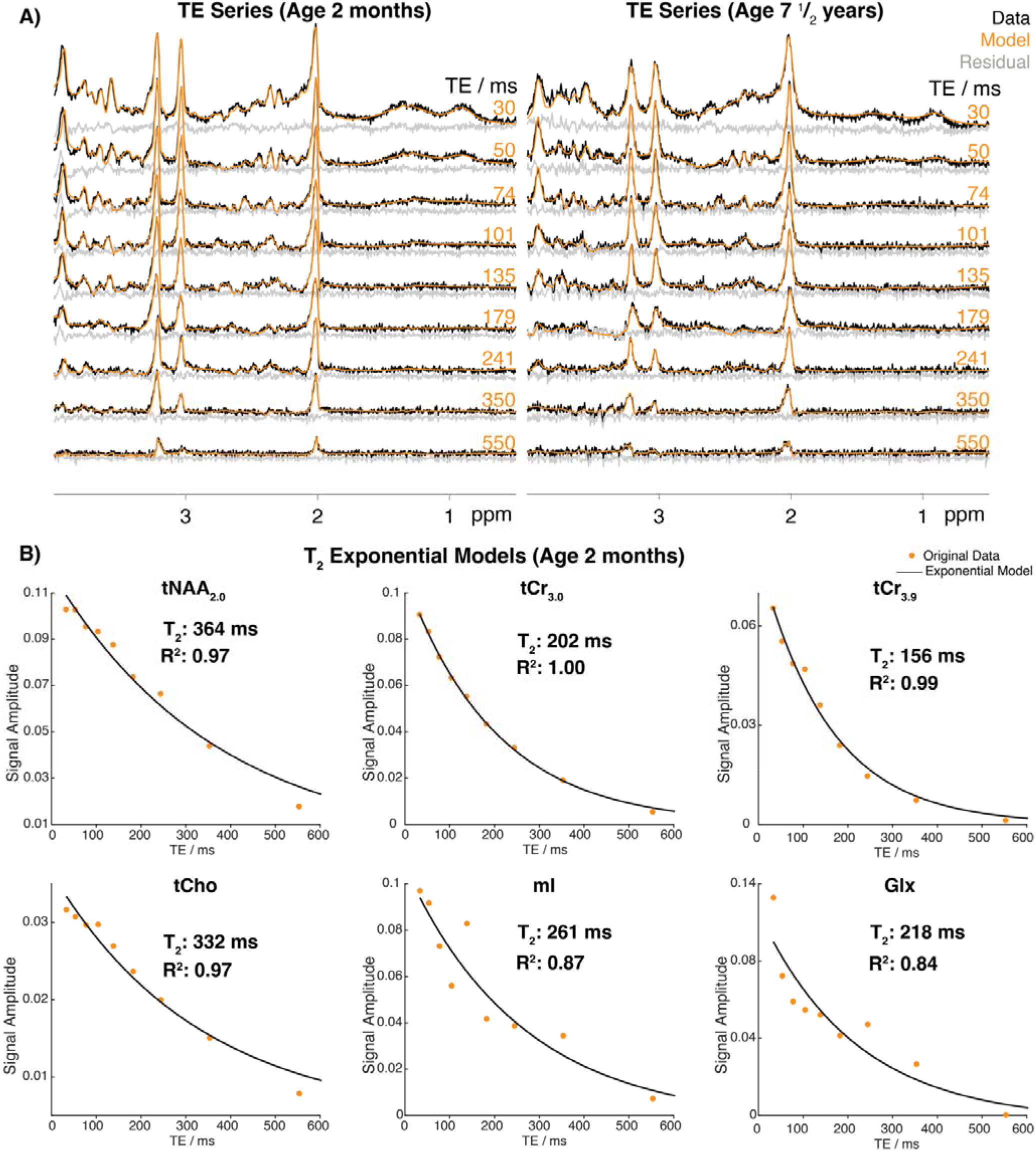
A) Example TE-series spectra, models, and fit residuals from a 2-month-old infant and a 90-month-old-child. B) TE exponential decay modelling to determine metabolite T_2_ relaxation times, using the modelled metabolite coefficients from the 2-month-old infant spectra in A.

T_1_ and T_2_ estimates are plotted against age in months in Figures 3 and 4 respectively, with linear models and Spearman correlation coefficients shown. The extrapolated intercepts of the linear age models provide metabolite T_1_ and T_2_ relaxation times in the thalamus at birth. The slope of the linear age models gives the rate of change of relaxation times per month up until 7 years of age. T_1_ and T_2_ relaxation times at any given age (in months) between birth and 95 months of age can be calculated for each metabolite from the linear model equations provided (Table 1).

**Figure 3:**
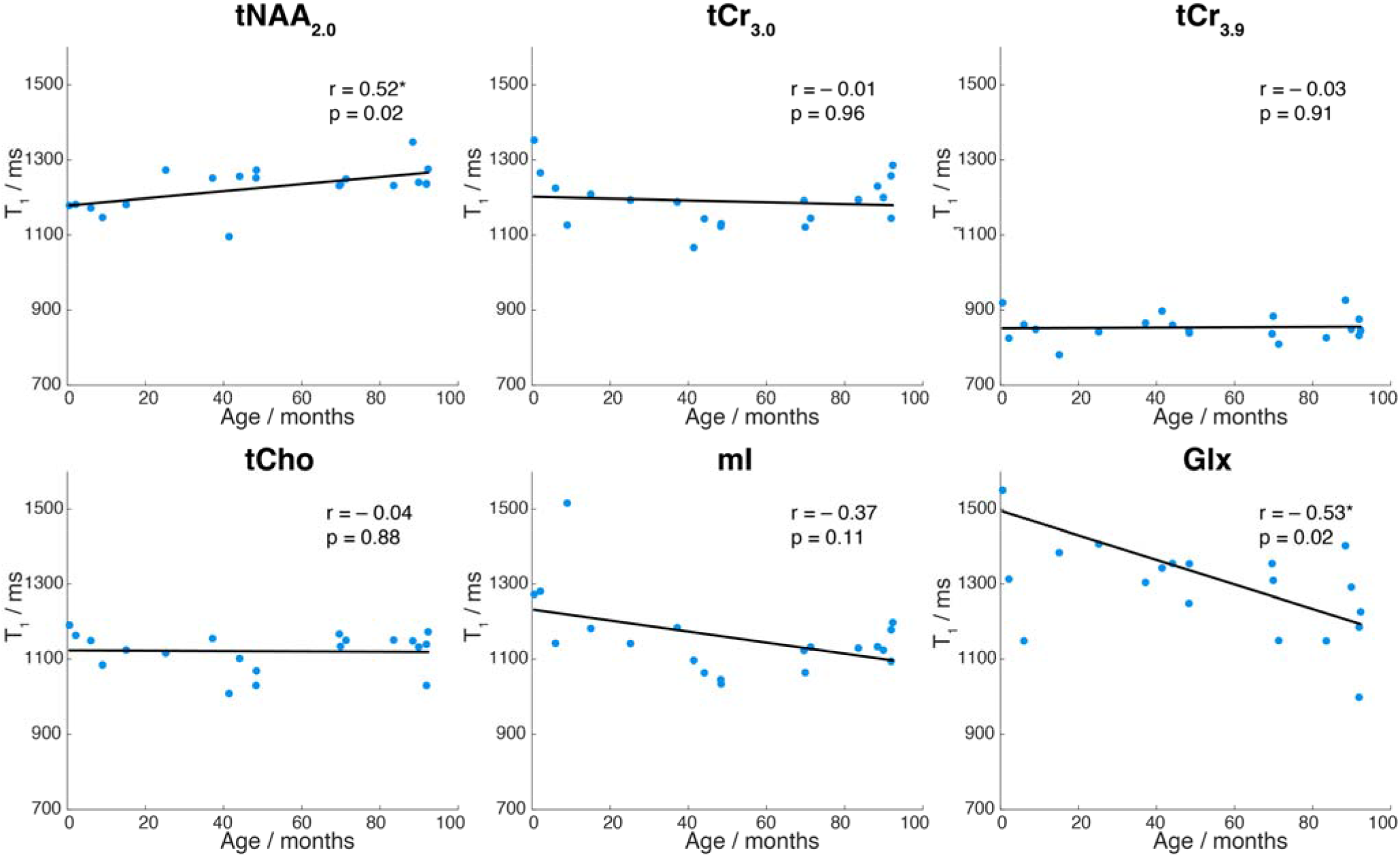
Correlations between T_1_ relaxation times and age in months, assessed by Spearman r and p values for each metabolite. Linear models are fitted to the datapoints, and the line is represented by T_1,Age_ = slope × Age + intercept for metabolites showing significant change in T_1_ with age (* indicates p < 0.05), as summarized in Table 1.

**Figure 4:**
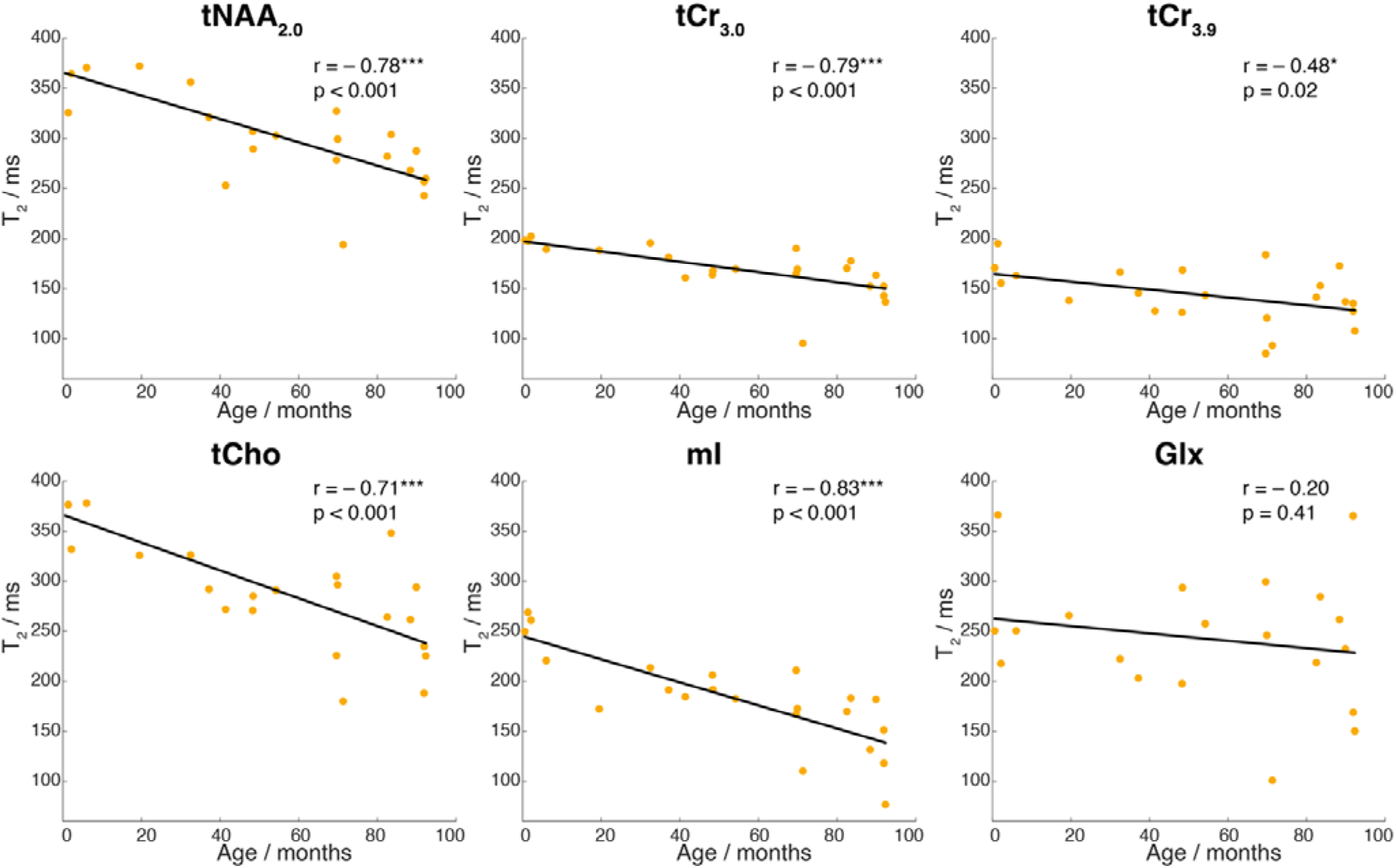
Correlations between T_2_ relaxation times and age in months, assessed by Spearman’s r and p values for each metabolite. Linear models are fitted to the datapoints, and the line is represented by T_2,Age_ = slope × age + intercept for metabolites showing significant change in T_2_ with age (* indicates p < 0.05, *** indicates p < 0.001), as summarized in Table 1.

## Discussion

In this work, we present new metabolite T_1_ and T_2_ relaxation reference values for pediatric MRS spanning 0–7 years of age. These pediatric T_1_ and T_2_ relaxation times differ substantively from published adult reference values^9^, including those currently used by default in nearly every MRS software. T_2_ relaxation times are generally higher than the adult reference values^10^, showing downward trends with age, while T_1_ showed a more variable pattern. Both sets of reference values are recommended for future pediatric quantification, because accurate quantification of concentrations relies upon relaxation correction with appropriate reference values.

The spectral modeling approach used here for the TI series has previously been applied to analogous recent work across the adult lifespan^11^, and similar challenges were observed in both datasets. Specifically, TIs that are close to metabolite null-points are the most challenging to model, and as before^11^ we used a more flexible spline baseline for TI 637 ms and no lineshape convolution for TI 637 and 780 ms. Although it is not ideal to change modelling parameters for spectra with a given series, this tailoring was essential for successful modelling of the dominant MM signals (and for avoiding metabolite lineshape estimation errors) close to metabolite null-points.

Modelling of TE series is generally more straightforward, because signal polarity is well-defined. However, the final TE-800 ms datapoint was difficult to model due to extremely low SNR. Therefore, we excluded this datapoint from the exponential modeling. Metabolite signals in TE 550 ms spectra (Figure 2) are sufficiently decayed to perform T_2_ exponential decay models to confidently estimate T_2_ relaxation times. Exponential decay curves were challenging to model for mI and Glx, likely due to J-evolution, and tCr_3.9_, due to overlap with the large residual water signal.

These results show that, in addition to being a dynamic period with changes in metabolite concentration^2,3,13,15,29^, early life also sees significant changes in metabolite relaxation. Both between the metabolites, and between T_1_ and T_2_, there is heterogeneity in these changes. This may reflect differential sampling of the changing biophysical microenvironment, or it may signal the need for further work. In contrast to our prior study in adults, ages 20 – 60+ years, where T_1_ relaxation times of most metabolites, including tNAA_2.0_ and Glx, decreased with age^11^, we found that T_1_ for NAA increased though age 7. This suggests that there is a point in later childhood or adolescence where T_1_ for NAA begins to decrease. Future studies are needed to establish this and whether it is consistent across all tissue types. Earlier studies at 1.5T^13,29^ also showed significant decreases in T_1_ of tCr_3.0_ within our age range in parieto-occipital regions; however, our contrasting findings suggest that there are either acquisition-dependent or tissue-specific differences in these pediatric relaxation parameters.

Decreases in T_2_ relaxation time with age, seen in our data and previous pediatric metabolite T_2_ studies at 1.5T^13^, are likely attributed to reduced water content and increased cellular viscosity^30^. Metabolite T_2_s continue to decrease throughout healthy adulthood^12^, for these reasons and due to increasing iron deposition^31,32^.

While the sample size of this study is not small and the reference values here are an improvement on the literature to-date, they raise enough questions to motivate larger studies including multiple regions and more extensive age sampling. Future analyses will probe not just the linear direction, but the shape of changes in relaxation times with age across different regions in the brain. The increased sample size will better power age modeling with improved anatomical and age coverage. It is likely that 2D modelling^33–35^ of the TI-/TE-series dataset, will prove to be a more SNR-efficient approach, using information from high-SNR spectra to guide modeling of TI-series spectra close to metabolite null-points and TE-series spectra at long echo times.

The scope of this technical note is to report new metabolite relaxation reference values. More in-depth analyses are required to quantitatively assess these values in the context of adult cohorts acquired with the same methodology^11,12^, and given the unsampled gap between cohorts (7-20 years), may have to wait for additional planned datasets. Using these age-appropriate relaxation parameters will change pediatric metabolite concentration estimates – both on average and potentially in influencing concentration-age relationships. Across the adult lifespan, T_1_ and T_2_ both tend to decrease with age^11,12^, effects that change metabolite concentration estimates^36^, but tend to counteract each other^37^ in influencing concentration-age correlations while including also the water relaxation times^38^. In contrast, corrections for tNAA_2.0_ based upon this cohort’s observations (of increasing T_1_ and decreasing T_2_ with age) would tend to accentuate each other in influencing age curves. Clearly, more data and further investigation are necessary to understand the impact of changing relaxation times with age on metabolite concentrations and their trajectories with age for pediatric populations.

## Conclusion

This work provides T_1_ and T_2_ relaxation times at 3T from birth to 7 years of age for the first time. Significant age-related changes in relaxation times are observed for multiple metabolites in the thalamus. Therefore, use of the linear equations provided here to estimate age-specific relaxation times is encouraged for pediatric MRS studies.

## Acknowledgements

This work was supported by National Institutes of Health (NIH) grants K99 HD118185, R01 EB032788, R01 EB016089, R01 EB023963, R01 EB035529, R21 EB033516, R00 AG080084, and P41 EB031771.

