## Supplementary Material for "Metabolite Relaxation Times at 3T From Birth to 7 Years"

**Summary MRSinMRS Report**

| **Table S1: Hardware** | |
| --- | --- |
| a. Field strength [T] | 3 T |
| b. Manufacturer | Philips |
| c. Model (software version if available) | R 5.7.1 |
| d. RF coils: nuclei (transmit/receive), number of channels, type, body part | 1H, 32 channel, Head |

| **Table S2:** **Acquisition** | |
| --- | --- |
| **Metabolite TI-Series** |  |
| a. Pulse sequence | Philips PRESS |
| b. Volume of Interest (VOI) | Bilateral thalamus |
| c. Nominal VOI size [mm^3^] | 23 × 30 × 23 |
| d. Repetition time (TR), echo time (TE),  inversion time (TI) [ms] | TR 2000 TE 30 TI 300, 400, 511, 637, 780, 947, 1148, 1400 |
| e. Total number of transients per spectrum | 24 per TI |
| f. Additional sequence parameters |  |
| g. Water suppression method | CHESS (140 Hz bandwidth) |
| h. Shimming method | 2^nd^ order pencil beam |
| i. Trigger or motion correction | No trigger or active motion correction |
| **Water TI-Series** |  |
| a. Pulse sequence | Philips PRESS |
| b. Volume of Interest (VOI) | Bilateral thalamus |
| c. Nominal VOI size [mm^3^] | 23 × 30 × 23 |
| d. Repetition time (TR), echo time (TE),  inversion time (TI) [ms] | TR 2000 TE 30 TI 300, 400, 511, 637, 780, 947, 1148, 1400 |
| e. Total number of transients per spectrum | 2 per TI |
| f. Additional sequence parameters |  |
| g. Water suppression method | None |
| h. Shimming method | 2^nd^ order pencil beam |
| i. Trigger or motion correction | No trigger or active motion correction |
| **Metabolite TE-Series** |  |
| a. Pulse sequence | Philips PRESS |
| b. Volume of Interest (VOI) | Bilateral thalamus |
| c. Nominal VOI size [mm^3^] | 30 (RL) × 23 × 23 |
| d. Repetition time (TR), echo time (TE) [ms] | TR 3000 TE 30, 50, 74, 101, 135, 179, 241, 350, 550 |
| e. Total number of transients per spectrum | 16 per TE |
| f. Additional sequence parameters |  |
| g. Water suppression method | VAPOR (100 Hz bandwidth) |
| h. Shimming method | 2^nd^ order pencil beam |
| i. Trigger or motion correction | No trigger or active motion correction |
| **Water TE-Series** |  |
| a. Pulse sequence | Philips PRESS |
| b. Volume of Interest (VOI) | Bilateral thalamus |
| c. Nominal VOI size [mm^3^] | 23 × 30 × 23 |
| d. Repetition time (TR), echo time (TE) [ms] | TR 2000 TE 30, 50, 74, 101, 135, 179, 241, 350, 550 |
| e. Total number of transients per spectrum | 2 per TE |
| f. Additional sequence parameters |  |
| g. Water suppression method |  |
| h. Shimming method | 2^nd^ order pencil beam |
| i. Trigger or motion correction | No trigger or active motion correction |

| **Table S3:** **Data Analysis Methods and Outputs** | |
| --- | --- |
| a. Analysis software | Osprey 2.5.0 |
| b. Processing steps deviating from Osprey | Manual phase correction of TI-series spectra |
| c. Output measure | Metabolite amplitude (from A_amplMets_Voxel_1_Basis.tsv) |
| d. Quantification references and assumptions, fitting model assumptions | Basis set list: Asc, Asp, CrCH_3_ at 3.0 ppm, CrCH_2_ at 3.9 ppm, GABA, GPC, GSH, Gln, Glu, Lac, mI, NAA_ace_ at 2.0 ppm, NAA_asp_, NAAG_ace_ at 2.0 ppm, NAAG_asp,glu_, PCh, PE, sI, Tau, MM09, MM12, MM14, MM17, MM20, Lip09, Lip13, Lip20 Fitting method: Osprey No convolution for TIs 637 and 780 ms Knot spacing:  Metabolite TI-Series 0.4 for TIs 300, 400, 511, 780, 947, 1148, 1400 0.2 for TIs 637 Metabolite TE-Series 0.4 for all TEs |

| **Table S4:** **Data Quality** | |
| --- | --- |
| **Metabolite TI-Series (TI: 1400 ms)** |  |
| a. SNR (tCr) | 43 ± 11 |
| b. Linewidth at FWHM (tCr) [Hz] | 7.5 ± 4.2 |
| c. Data exclusion criteria | tCr linewidth > 12 Hz, tCr SNR < 5, Lipid contamination |
| **Metabolite TE-Series (TE: 30 ms)** |  |
| a. SNR (Cr) | 39 ± 11 |
| b. Linewidth at FWHM (Cr) [Hz] | 6.2 ± 1.5 |
| c. Data exclusion criteria | tCr linewidth > 12 Hz, tCr SNR < 5, Lipid contamination |
